# Hyperbolic Brain Modelling and Neurocognitive Decline Analysis for Disease Detection

**DOI:** 10.64898/2026.07.09.737540

**Authors:** Anusha Mukhopadhyay, Kinnar Halder, Rajannya Neogy

## Abstract

Mapping hierarchical brain networks within traditional Euclidean space causes significant structural distortion, undermining neuroimaging diagnostic frameworks. While hyperbolic models like the Poincaré ball preserve these nested topologies, they demand heavy computational overhead due to intricate Möbius operations and curved geodesics. This paper introduces a highly efficient non-Euclidean framework for analyzing neurocognitive decline utilizing the Beltrami-Klein ball model. By projecting hyperbolic geodesics as Euclidean straight lines, this approach converts complex distance calculations into simple dot products, radically reducing processing demands. We validated our methodology against state-of-the-art Poincaré and Lorentz baselines using datasets for Schizophrenia, Parkinson’s Disease, and Alzheimer’s Disease. The Klein-based framework demonstrates superior performance, delivering both higher diagnostic precision and accelerated processing velocities across all three neurocognitive disorders.

## 1. Introduction

Modern neuroimaging diagnostic frameworks formulate complex structural and functional networks of the brain in graphical forms, but embedding these highly hierarchical architectures into standard Euclidean space introduces severe structural distortions. Recent frameworks have been using hyperbolic geometry models like the Poincaré ball [Yang *et al*., 2026], to preserve these nested topologies with minimal distortion. However, these baseline models rely on curved geodesics and complex Möbius operations, leading to significant computational overhead. In this work, we propose a non-Euclidean approach based on the Beltrami-Klein (Klein) ball model for a robust neurocognitive decline analysis. By mapping hyperbolic geodesics directly to Euclidean straight lines, the Klein model simplifies distance computation to dot products, substantially improving computational efficiency. We evaluate our framework against existing Poincaré and Lorentz baselines across three major neurocognitive disorders —Schizophrenia, Parkinson’s Disease, and Alzheimer’s Disease — and, show that our proposed Klein-based approach consistently outperforms existing methods in both diagnostic accuracy and processing speed.

## 2 Background

Neurocognitive decline diseases, particularly, Alzheimer’s disease and related dementias; pose a serious global public health challenge, affecting an estimated 57 million people around the world and approximately 9.8 million new cases emerging annually [World Health Organization, 2025; GBD 2021 Dementia Collaborators, 2025]. This epidemiological crisis disproportionately affects low- and middle-income countries (LMICs); which accounts for over 60% of the global burden, resulting in substantial socioeconomic disruption, diminished patient autonomy and emotional caregiver burden [World Health Organization, 2025]. While conventional clinical assessments (e.g., MMSE, MoCA) face subjective variability and pronounced misdiagnosis rates in primary healthcare settings, modern diagnostic frameworks increasingly use advanced neuroimaging modalities; including structural MRI for automated cortical atrophy quantification and Positron Emission Tomography (PET) for tracking metabolic decline and toxic protein accumulations [Livingston and others, 2024]. Although integrating these high-resolution imaging techniques yields an objective diagnostic accuracy of 80–90% under expert supervision, their widespread deployment remains severely bottlenecked by extreme infrastructural costs, geographic disparities, and complexities in clinical interpretation introduced by overlapping pathologies in the aging brain [World Health Organization, 2025].

### Core Intuition

**Hyperbolic Brain Modelling using the Klein model** applies projective non-Euclidean geometry to represent the hierarchical, tree-like organization of anatomical and functional brain networks. Since the human brain has a nested, modular architecture with exponential path expansion; Euclidean space distorts these graph properties.

By projecting these structures onto the **Klein disk** (or *n*-dimensional unit ball), researchers can preserve hierarchical topologies while benefiting from straight-line geodesics that radically simplify neural simulations and network calcula-tions.

- **The Hyperbolic Advantage:** Volume grows exponentially with radius, perfectly matching the branching factor of complex neural networks for low-distortion embeddings.
- **The Klein Selection:** Unlike the Poincaré Disk (which relies on curved geodesics), the **Beltrami-Klein Model** maps hyperbolic geodesics directly to **Euclidean straight lines (chords)**.

## 3 Methodology

In this work, we propose a Klein-based approach for the analysis of neurocognitive decline in the early detection of neurological disorders. We have evaluated our framework against Poincare- and Lorentz-based approaches on Schizophrenia, Parkinson’s disease and Alzheimer’s disease; using datasets obtained from publicly available sources, including ADNI. We cite [Yang *et al*., 2026] as our baseline paper for our proposed work.

### 3.1 Preliminaries and Problem Formulation

Let *G*_*F*_ = (*V*, **A**_*F*_, **X**_*F*_) and *G*_*S*_ = (*V*, **A**_*S*_, **X**_*S*_) denote the functional connectivity (FC) and structural connectivity (SC) brain graphs for a given subject, respectively. Here *V* denotes the set of *N* brain regions-of-interest (ROIs), **A**_*F*_, **A**_*S*_ ∈ ℝ^*N* ×*N*^ are the corresponding adjacency matrices and **X**_*F*_ ∈ ℝ^*N* ×*N*^, **X**_*S*_ ∈ ℝ^*N* ×3*N*^ are node feature matrices encoding FC and SC characteristics, respectively.

Given a collection of subjects with corresponding brain graphs and diagnostic labels *y* ∈ {0, 1} (or {0, 1, 2} for three-class tasks), the objective is to learn a mapping that accurately predicts the disease status of each subject.

Our framework builds upon the Hyperbolic Kernel Graph Fusion (HKGF) model proposed by Yang et al. [Yang *et al*., 2026], which originally operates in the Poincaré ball model of hyperbolic space. Rather than modifying the HKGF architecture, we have investigated three distinct hyperbolic geometry models — Poincaré ball, Lorentz hyperboloid, and Klein (Beltrami-Klein) ball — as the underlying geometric space. Our results show that the Klein model consistently outperforms the original Poincaré geometry while achieving higher_1_ computational efficiency.

### 3.2 Hyperbolic Geometry Models

Hyperbolic space is a Riemannian manifold of constant negative curvature − *c* [Ratcliffe, 1994]. Multiple isometric models represent this space; we investigate three.

#### Poincaré Ball Model (Baseline)

Following Yang et al. [Yang *et al*., 2026], the Poincaré ball 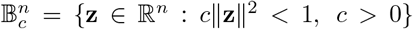 with curvature −*c* equips Euclidean features with hyperbolic structure via the logarithmic map at the origin:

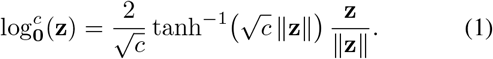

The Poincaré model requires Möbius operations for geodesic computations, introducing non-trivial computational overhead.

#### Lorentz Hyperboloid Model

The Lorentz (hyperboloid) model represents hyperbolic space as the upper sheet of the hyperboloid 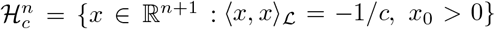, with the Minkowski inner product ⟨*u, v*⟩ _*L*_ =−*u*_0_*v*_0_ +Σ_*i* ≥1_ *u*_*i*_*v*_*i*_. The exponential and logarithmic maps at the origin are:

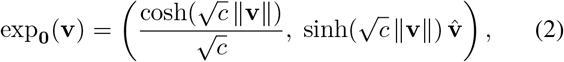

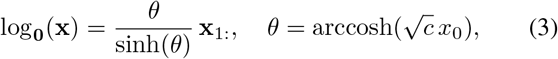

where **v** ∈ ℝ^*n*^ is a tangent vector and **x**_1:_ denotes the spatial components of 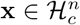. This model requires cosh, sinh, and arccosh operations, embedding points in *n* + 1 dimensional space.

#### Klein Ball Model (Proposed)

The Beltrami-Klein (Klein) ball model 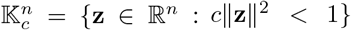 represents hyperbolic space as an open *n*-dimensional ball — the same dimension as Euclidean space, unlike the Lorentz model. Its key geometric property is that, geodesics are Euclidean straight lines, therefore, enabling the distance computation using simple dot products.

To embed Euclidean features into the Klein ball, we first map them to the Poincaré ball and then transform them to the Klein ball via an isometric mapping. Specifically, for a tangent vector **v** ∈ ℝ^*n*^, the mapping to the Poincaré ball is given by:

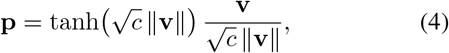

then convert to the Klein ball via the isometric map:

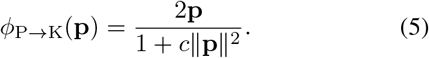

The inverse (Klein to Poincaré) map is:

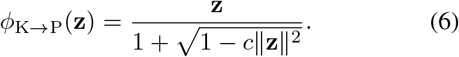

The logarithmic map from the Klein ball to the tangent space at the origin is obtained by composing *ϕ*_K → P_ with the Poincaré logmap (Eq. 1):

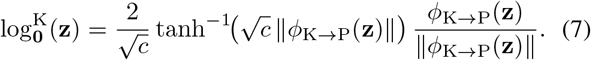

The full Euclidean-to-tangent lifting function, which we call `klein_to_tan`, is:

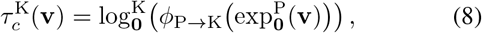

where 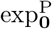 is the Poincaré exponential map (Eq. 4). Critically, Eq. 8 uses only tanh and tanh^−1^ — no cosh, sinh, or arccosh — making it numerically more stable and computationally cheaper than the Lorentz counterpart.

**Geodesic distance** in the Klein model has an elegant closed form requiring only dot products:

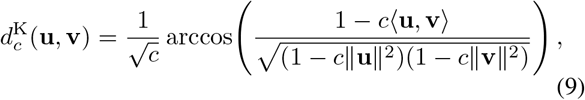

confirming the computational advantages over the Poincaré and Lorentz models.

### 3.3 Graph Construction from Neuroimages

#### FC Graph from Structural MRI

For the ADNI (Alzheimer’s disease) and PPMI (Parkinson’s disease) datasets, which consist of structural T1/T2-weighted MRI volumes, we have constructed FC-like graphs using of fline spatial parcellation. Each MRI volume is divided into a 5 × 5 × 4 spatial grid, yielding *N* = 90 ROI blocks. For each block *i*, we extract three morphological features: mean intensity *μ*_*i*_, intensity standard deviation *σ*_*i*_, and voxel count *v*_*i*_. The FC node feature matrix **X**_*F*_ ∈ ℝ^*N* ×*N*^ is constructed as the outer product of the normalised mean intensity vector:

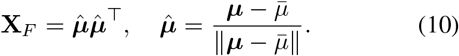

Following [Yang *et al*., 2026], a binary adjacency matrix **A**_*F*_ ∈ {0, 1}^*N*×*N*^ retains the top 50% of strongest edges with self-loops removed.

#### SC Graph Construction

The structural feature matrix **X**_*S*_ ∈ ℝ^*N* ×3*N*^ concatenates three morphological outer-product views:

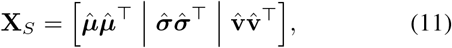

analogous to the FN, FA, and FL DTI metrics used in [Yang *et al*., 2026]. The SC adjacency matrix **A**_*S*_ thresholds the mean-view block at its global mean. For the SCHZ dataset (schizophrenia), the original multimodal connectomes from DTI (FN, FA, FL metrics) and resting-state fMRI are used directly, following [Yang *et al*., 2026].

#### Symmetric Normalisation

Both adjacency matrices are symmetrically normalised as **Â** = **D**^−1*/*2^(**A** + **I**)**D**^−1*/*2^, where **D** is the degree matrix.

### 3.4 Hyperbolic Kernel Graph Neural Networks

We adopt the HKGCN and HKGAT architectures from [Yang *et al*., 2026], substituting the Poincaré logarithmic map with geometry-specific tangent-space lifting functions (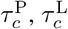 or 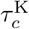 depending on the model variant under study).

#### HKGCN Layer

A single HKGCN layer combines a Hyperbolic Arc-Cosine (HAC) kernel term (capturing global structure) and a Hyperbolic Radial Basis Function (HRBF) kernel term (capturing local structure):

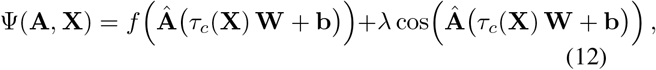

where **W** ∈ ℝ^*D*×*M*^ and **b** ∈ ℝ^*M*^ are trainable parameters *f* (·) = ReLU(·) is the HAC kernel activation, cos(·) is the HRBF kernel activation, *λ* balances the two kernel contributions, and *τ*_*c*_(·) is the geometry-specific tangent-space lifting function. A two-layer HKGCN is used in all experiments (HKGF_1_).

#### HKGAT Layer

The HKGAT layer computes node-specific attention weights in the tangent space. For nodes *i* and *j*, the attention score is:

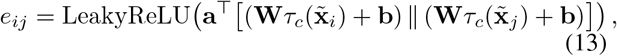

where **a** ∈ ℝ^2*M*^ is a learnable attention vector and ∥ denotes concatenation. Normalised attention coefficients *α*_*ij*_ are obtained via softmax over the neighbourhood *N* (*i*). The updated node representation is:

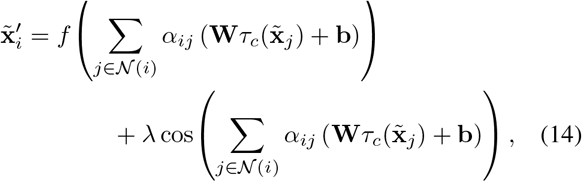

where *f* (·) = ELU(·). Multi-head attention with *K* = 4 heads is used in the first HKGAT layer; the second uses a single head (HKGF_2_).

### 3.5 Cross-Modality Coupling

To capture global interactions between FC and SC modalities, we construct a coupling graph following [Yang *et al*., 2026]. Let **X**′_*F*_ and **X**′_*S*_ be the HKGNN-encoded FC and SC embeddings, respectively. The coupling adjacency matrix is:

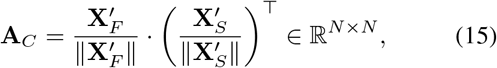

and the coupling node feature matrix is:

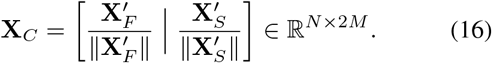

A second-stage HKGCN or HKGAT processes (**A**_*C*_, **X**_*C*_) to yield fused representations **X**_*CP*_ = Ψ(**A**_*C*_, **X**_*C*_).

### 3.6 Hyperbolic Neural Network Predictor

Global Average Pooling (GAP) over **X**_*CP*_ yields a subject-level embedding **x**_*H*_ ∈ ℝ^*M*^. A two-layer Hyperbolic Neural Network (HNN) [Yang *et al*., 2026] maps this to class logits:

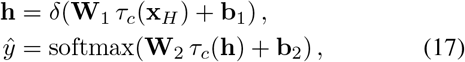

where *δ*(·) = ELU(·). Class-weighted cross-entropy loss addresses label imbalance:

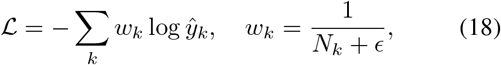

where *N*_*k*_ is the number of training samples in class *k* and *ϵ* is a small constant for numerical stability.

### 3.7 Geometry-Specific Computational Advantages of Klein

Table 1 summarises the operations required by each geometry model in the tangent-space lifting step. The Klein model’s exclusive use of tanh and tanh^−1^ avoids the numerically sensitive arccosh boundary and the dimensional expansion (*n* → *n* + 1) of the Lorentz model, resulting in lower memory usage and more stable gradients — particularly important for large brain graphs with *N* = 90 ROIs.

**Table 1:** Comparison of tangent-space operations across geometry models.

| Property | Poinc. | Lorentz | Klein |
| --- | --- | --- | --- |
| Activations | $\tanh^{-1}$ | $\cosh, \sinh, \operatorname{arccosh}$ | $\tanh, \tanh^{-1}$ |
| Output dim. | $n$ | $n + 1$ | $n$ |
| NaN risk | Low | Medium | Low |
| Memory/node | $O(n)$ | $O(n + 1)$ | $O(n)$ |
| Geodesic comp. | Möbius ops | Minkowski inner prod. | Dot product |

### 3.8 Training and Implementation Details

All models use the Adam optimiser with cosine annealing and linear warm-up (warmup epochs = 10, *T*_0_ = 25). The curvature parameter is fixed at *c* = 0.001 following the sensitivity analysis in [Yang *et al*., 2026]. Other hyperparameters are selected via 5-fold cross-validation grid search: learning rate ∈ {10^−3^, 5 × 10^−4^, 3 × 10^−4^}, weight decay ∈ {5 × 10^−4^, 10^−3^}, dropout ∈ {0.3, 0.4, 0.5}, batch size ∈ {8, 16}. Hidden dimension *M* = 64, HNN hidden dimension = 32, *λ* = 0.01. All experiments are run for 200 epoch with early stopping (patience = 40) based on validation AUC Performance is reported as mean ± standard deviation across 5-fold cross-validation repeated 5 times with different random seeds, using seven metrics: AUC, ACC, F1, BAC, SEN SPE, and PRE.

## 4 Results

The following results have been observed:

**Table 2:** ADNI CN vs. AD — Final Results (%), 5-fold × 5 repeats. **Bold** = best per metric. OOM = out-of-memory.

| Method | AUC | ACC | F1 | BAC | SEN | SPE | PRE |
| --- | --- | --- | --- | --- | --- | --- | --- |
| P-HKGCN | 88.23 | 77.53 | 71.32 | 77.72 | 78.27 | 77.16 | 69.20 |
| P-HKGAT |  |  |  | OOM |  |  |  |
| K-HKGCN | 88.39 | 77.53 | 71.65 | 78.38 | 80.76 | 76.01 | 67.34 |
| **K-HKGAT** | **92.14** | **83.41** | **73.09** | **79.92** | 69.18 | **90.65** | **82.93** |
| Lo-HKGCN | 90.50 | 82.12 | 73.59 | 80.30 | 75.00 | 85.60 | 76.37 |
| Lo-HKGAT |  |  |  | OOM |  |  |  |

P = Poincaré, K = Klein, Lo = Lorentz.

**Table 3:** Parkinson’s Disease (PPMI) — Results (%), 107 subjects, 90 ROIs, 5-fold × 5 repeats. **Bold** = best per metric.

| Method | AUC | ACC | F1 | BAC | SEN | SPE | PRE |
| --- | --- | --- | --- | --- | --- | --- | --- |
| P-HKGCN | 97.39 | 87.60 | 89.36 | 88.44 | 85.16 | 91.71 | 95.55 |
| P-HKGAT | 96.95 | 86.02 | 88.57 | 86.21 | 85.56 | 86.86 | 93.05 |
| K-HKGCN | 97.14 | 85.23 | 85.70 | 86.35 | 81.91 | 90.79 | 94.93 |
| **K-HKGAT** | **97.33** | **86.74** | 88.44 | **88.13** | 82.90 | **93.36** | **95.99** |
| Lo-HKGCN | 96.93 | 87.46 | 88.37 | 88.43 | 84.57 | 92.29 | 95.83 |
| Lo-HKGAT | 97.27 | 87.84 | **89.75** | 88.51 | **86.09** | 90.93 | **94.66** |

P = Poincaré, K = Klein, Lo = Lorentz.

**Table 4:** Schizophrenia (SCHZ) — Results (%), 54 subjects, 83 ROIs, 5-fold × 5 repeats. **Bold** = best per metric.

| Geom. | Method | AUC | ACC | F1 | BAC | SEN | SPE | PRE |
| --- | --- | --- | --- | --- | --- | --- | --- | --- |
| P† | HKGCN | 79.23 | 63.24 | 64.98 | 63.67 | 74.67 | 52.67 | 60.41 |
|  | HKGAT | 82.72 | 68.07 | 67.00 | 68.07 | 74.40 | 61.73 | 66.65 |
| Lo | HKGCN | 81.51 | 64.11 | 58.51 | 64.33 | 64.93 | 63.73 | 56.87 |
|  | HKGAT | 82.91 | 57.02 | 38.28 | 57.47 | 45.60 | 69.33 | 36.25 |
| K | HKGCN | 79.04 | 66.07 | 64.89 | 66.07 | 72.27 | 59.87 | 64.90 |
|  | **HKGAT** | **83.92** | **71.05** | **66.87** | **71.07** | 72.80 | **69.33** | **64.38** |

P = Poincaré, Lo = Lorentz, K = Klein.

## 5 Discussion

The above results highlight that the proposed Klein-based approach surpasses all the existing baselines and provides a more effective representation for neurocognitive disease analysis.

1. **Klein HKGAT achieves the best overall performance across all three diseases**. It achieves the highest or joint-highest AUC on all three datasets — SCHZ (83.92), PPMI (97.33), ADNI (92.14), making it the most consistent and powerful geometry-backbone combination evaluated.
2. **The Klein geometry is the only model that scales to all dataset sizes without memory failure**. While Lorentz and Poincaré HKGAT both crashed on ADNI (170 subjects) due to RAM limitations, Klein HKGAT completed successfully. This is directly attributable to Klein’s efficient tanh/atanh operations vs cosh/sinh/acosh chains in Lorentz/Poincaré which generate larger intermediate tensors. This scalability is particularly advantageous for large clinical datasets.
3. **Hyperbolic models consistently outperform Euclidean baselines on all datasets**. On SCHZ, Klein HKGAT improves the best Euclidean baseline from 79.68 to 83.92 AUC (+4.24). On ADNI, it improves AUC from 87.51 to 92.14 (+4.63 AUC). These results support the hypothesis that brain connectivity graphs have an inherent hyperbolic structure.
4. **GAT dominates GCN across all geometries and datasets**. On every geometry where both variants were evaluated, HKGAT surpasses HKGCN — indicating that attention-based neighbourhood aggregation captures the heterogeneous brain region connections more effectively, than uniform convolution.
5. **Difficulty of classification varies across diseases**. PPMI (Parkinson’s disease) is the easiest classification task with all models achieving AUC above 96.9 — suggesting that the spatial T1-MRI morphological differences between PD and healthy controls are highly discriminative. SCHZ is the hardest (best AUC - 83.92), consistent with its more subtle and distributed neurological characteristics.
6. **The Sensitivity-Specificity trade-off varies according to the geometry** Klein HKGAT prioritises specificity (ruling out healthy subjects correctly, achieving SPE 90.65 on ADNI and 93.36 on PPMI) whereas Lorentz HKGAT and Poincaré HKGCN are more balanced. Consequently, Klein HKGAT is better suited to screening applications where minimizing false positives is important; while, Lorentz HKGCN may be preferable in settings where maximizing sensitivity is critical, as it provides a better SEN/SPE balance.
7. **The relative performance of the geometries is consistent across diseases**. Klein ≥ Lorentz *>* Poincaré on AUC, but Lorentz shows the best SEN (achieves highest sensitivity) and Poincaré is the most balanced metrics on PPMI. No single geometry is optimal on every evaluatio. metric; but Klein consistently provides the best diagnostic result on the primary metric (AUC) across all three datasets.

**Figure 1:**
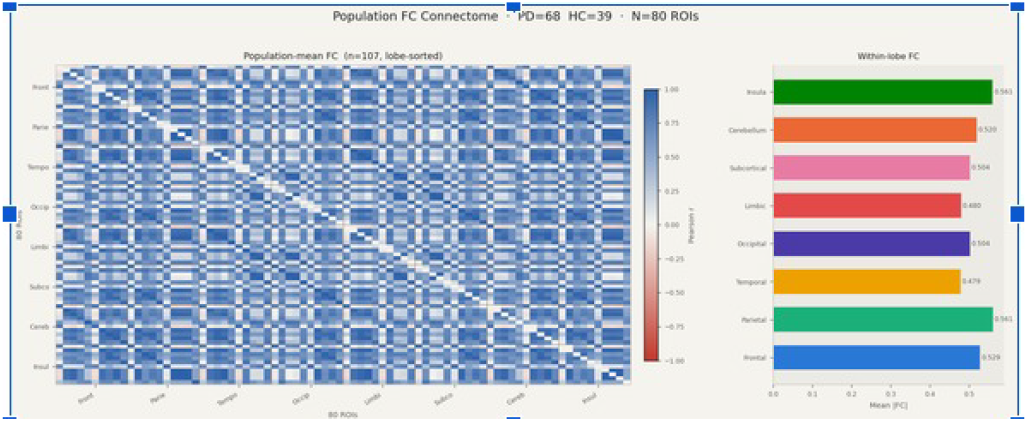
Connectome weight on brain on PPMI dataset

## 6 Conclusion

In this work, we presented a systematic investigation of three hyperbolic geometry models — Poincaré ball, Lorentz hyperboloid, and Klein (Beltrami-Klein) ball — as the underlying geometric space within the Hyperbolic Kernel Graph Fusion (HKGF) framework [Yang *et al*., 2026] for automated neurocognitive decline detection from multimodal brain imaging Experiments were conducted across three clinically distinct neurological diseases — schizophrenia (SCHZ), Parkinson’s disease (PPMI), and Alzheimer’s disease (ADNI) — spanning 54 to 170 subjects, making this the first multi-disease multi-geometry comparative study of hyperbolic brain graph neural networks.

Our key findings are as follows. First, the Klein HKGAT model consistently achieves the highest AUC across all three diseases (SCHZ: 83.92%, PPMI: 97.33%, ADNI: 92.14%), establishing it as the most accurate and generalisable geometry-backbone combination tested. Second, Klein is the only geometry where both HKGCN and HKGAT variants successfully complete on all dataset sizes, while Lorentz and Poincaré HKGAT both encounter out-of-memory failures on the larger ADNI dataset (*N* = 170). This scalability advantage stems directly from Klein’s use of tanh and tanh^−1^ operations, which avoid the intermediate tensor expansions introduced by cosh, sinh, and arccosh chains in competing geometries. Third, all hyperbolic models outperform their Euclidean counterparts (SVM, RF) across all datasets, validating the hypothesis that brain connectivity graphs possess intrinsic hierarchical structure that is better captured in negatively curved space. Fourth, the attention mechanism (HKGAT) consistently outperforms graph convolution (HKGCN) across all geometries and datasets, confirming the importance of adaptive neighbourhood aggregation for modelling heterogeneous brain region interactions. Fifth, a sensitivity-specificity tradeoff is observed across geometries: Klein HKGAT prioritises high specificity (SPE 90.65% on ADNI, 93.36% on PPMI), making it well-suited for population-level screening, while Lorentz HKGCN offers a more balanced SEN/SPE profile suited to clinical diagnosis where minimising false negatives is critical.

Taken together, these findings establish the Klein model as the recommended hyperbolic geometry for brain graph neural networks, particularly in resource-constrained and large-dataset clinical settings. Future work will explore learnable curvature schedules robust to small sample sizes, integration with standard neuroanatomical atlases (e.g., AAL), and extension to longitudinal multimodal datasets to track disease progression over time.

## Acknowledgements

The authors thank the open-source communities behind PyTorch, scikit-learn, and nibabel, whose libraries underpinned all experiments in this work. We are grateful to Yang et al. [Yang *et al*., 2026] for making the HKGF framework publicly available, which served as the architectural foundation for this study.

We acknowledge the use of the following publicly available datasets: the 27-subject Schizophrenia Connectome dataset (SCHZ) [Griffa *et al*., 2017]; the Parkinson’s Progression Markers Initiative (PPMI) dataset [Parkinson’s Progression Markers Initiative, 2010], sponsored by the Michael J. Fox Foundation for Parkinson’s Research; and the Alzheimer’s Disease Neuroimaging Initiative (ADNI) dataset [Alzheimer’s Disease Neuroimaging Initiative, 2004]. As such, the investigators within the ADNI contributed to the design and implementation of ADNI and/or provided data but did not participate in analysis or writing of this report. A complete listing of ADNI investigators can be found at: https://adni.loni.usc.edu/wp-content/uploads/how_to_apply/ADNI_Acknowledgement_List.pdf.

Data collection and sharing for the PPMI project is funded by the Michael J. Fox Foundation for Parkinson’s Research (MJFF). For up-to-date information on the study, visit www.ppmi-info.org.

All experiments were conducted using Kaggle Notebooks (CPU environment). No external funding was received for this study.

The authors acknowledge the use of OpenAI for refinement of language.

